# Alveolar macrophages play a key role in tolerance to ozone

**DOI:** 10.1101/2024.02.18.580749

**Authors:** Gregory J. Smith, Morgan Nalesnik, Robert M. Immormino, Jeremy M. Simon, Jack R. Harkema, Jason R. Mock, Timothy P. Moran, Samir N. P. Kelada

## Abstract

Acute exposure to ozone (O_3_) causes pulmonary inflammation and injury in humans and animal models. In rodents, acute O_3_-induced inflammation and injury can be mitigated by pre-exposure to relatively low concentration O_3_, a phenomenon referred to as tolerance. While tolerance was described long ago, the underlying mechanisms are not known, though upregulation of antioxidants has been proposed. To identify new mechanisms for O_3_ tolerance, we generated a mouse model in which female C57BL6/NJ mice were pre-exposed to filtered air (FA) or 0.8 ppm O_3_ for four days (4 hours/day), then challenged with 2 ppm O_3_ (3 hours) 2 days later, and phenotyped for airway inflammation and injury 6 or 24 hours thereafter. As expected, pre-exposure to O_3_ resulted in significantly reduced airway inflammation and injury at 24 hours, as well as reduced induction of antioxidant genes. Like previous studies in rats, tolerance was associated with changes in the frequency and proliferation of alveolar epithelial cells, but was not associated with upregulation of antioxidants, CCSP (SCGB1A1), or mucus. We found that alveolar macrophages (AMs) play a critical role in tolerance, as depletion of AMs using clodronate in mice pre-exposed to O_3_ restored many responses to acute O_3_ challenge. Further, AMs of O_3_ tolerized mice exhibited decreased expression of genes involved in cellular signaling via Toll-like receptors, MYD88, and NF-kB, and proinflammatory cytokine production. We conclude that O_3_ tolerance is highly, but not exclusively, dependent on AMs, and that further studies investigating how repeated O_3_ exposure induces hypo-responsiveness in AMs are warranted.

## INTRODUCTION

Exposure to the ambient air pollutant ozone is a pressing public health concern due to its association with cardiopulmonary morbidity and mortality^1,2^. It is well established that a single (acute) exposure to ozone causes airway inflammation, injury, and decreased lung function in humans^3–7^. Paradoxically, however, responses decrease–rather than increase–after repeated ozone exposures^8–17^, a process referred to as adaptation. Attenuated inflammation and injury have also been observed in rodent models^18,19^. Perhaps more strikingly, studies have shown that low-level exposures to ozone can protect rodents from subsequent challenges with higher, even lethal, concentrations of ozone^20–23^, a phenomenon referred to as tolerance^24^.

Several mechanisms underlying adaptation and/or tolerance to ozone have been proposed. These include upregulation of antioxidants to protect against free radicals and other oxidant chemical species^18,25^, which can vary by location the lung^26^. Upregulation of CCSP (SCGB1A1), which may have antioxidant properties^27^, has also been reported^28^. Likewise, increased mucus production has been posited as a potential protective mechanism because it contains antioxidants^29,30^, but this has not been evaluated further. Studies in rodent models have reported that exposure to ozone causes changes in the cellular composition of both the airways^30,31^ and alveoli^21^, which could alter subsequent responses to ozone. In long exposure duration models, the effects on the airway include increased frequency of mucin-producing epithelial cells and concomitant decreased ciliated cell frequency^31,32^. In models of tolerance per se, key findings include increased numbers of proliferating alveolar type 2 (AT2) cells^21,23^ and decreased alveolar surface area occupied by alveolar type (AT1) cells^21^, likely reflecting the lung’s compensatory response to the first occurrence of ozone-induced alveolar injury.

In reviewing the literature on ozone tolerance, we found that most, if not all, studies to date focused on the role of epithelial cell populations in the lung, leaving the role of other cell types, including leukocyte populations, unexamined. Additionally, almost all prior studies utilized rats^21–23,25^. Here, we developed a mouse model of ozone tolerance to enable us to evaluate previous hypotheses about mechanisms underlying tolerance, and also included an assessment of ozone dosimetry. We then evaluated the role of one leukocyte population, alveolar macrophages (AMs), in the development of tolerance, using both functional and genomic approaches. AMs are well-known contributors to acute ozone response^33–41^, and we and others have shown that ozone exposure alters the transcriptome of AMs^37,38,42^. Macrophage populations have been shown to play key roles in adaptation/tolerance to other stimuli (e.g., LPS and chlorine gas)^43,44^, and in these models macrophages exhibit marked transcriptional changes. Thus, we hypothesized that AMs may be key players in ozone tolerance and that tolerance stems from changes at the level of the AM transcriptome.

## METHODS

### Mice

Multiple studies have reported differences in ozone response by sex^45–49^. The results vary by time point, end point evaluated, and other factors. To limit the impact of sex differences on responses in our model of tolerance, we chose to focus on female mice, which have been observed to exhibit stronger inflammatory responses to ozone^45–47^. Female C57BL/6NJ mice (strain # 005304) were purchased from the Jackson Laboratory and housed in groups of ∼3-5/cage under normal 12-hour light/dark cycles with *ad libitum* food (Envigo 2929) and water on ALPHA-Dri bedding (Shepard). All mice used were in the age range of 8-11 weeks. All studies were conducted in compliance with University of North Carolina Institutional Animal Care and Use Committee guidelines in animal facilities approved and accredited by AAALAC.

### Ozone Exposures

Mice were exposed to filtered air (FA), 0.8 ppm ozone, or 2 ppm over the course of 7 days and harvested on day 8, as depicted in Figure 1A, using a computer-controlled, extremely precise ozone exposure system^50^. In subsequent experiments, mice were exposed to FA or 0.8 ppm ozone for four days and harvested on day 7.

**Figure 1.**
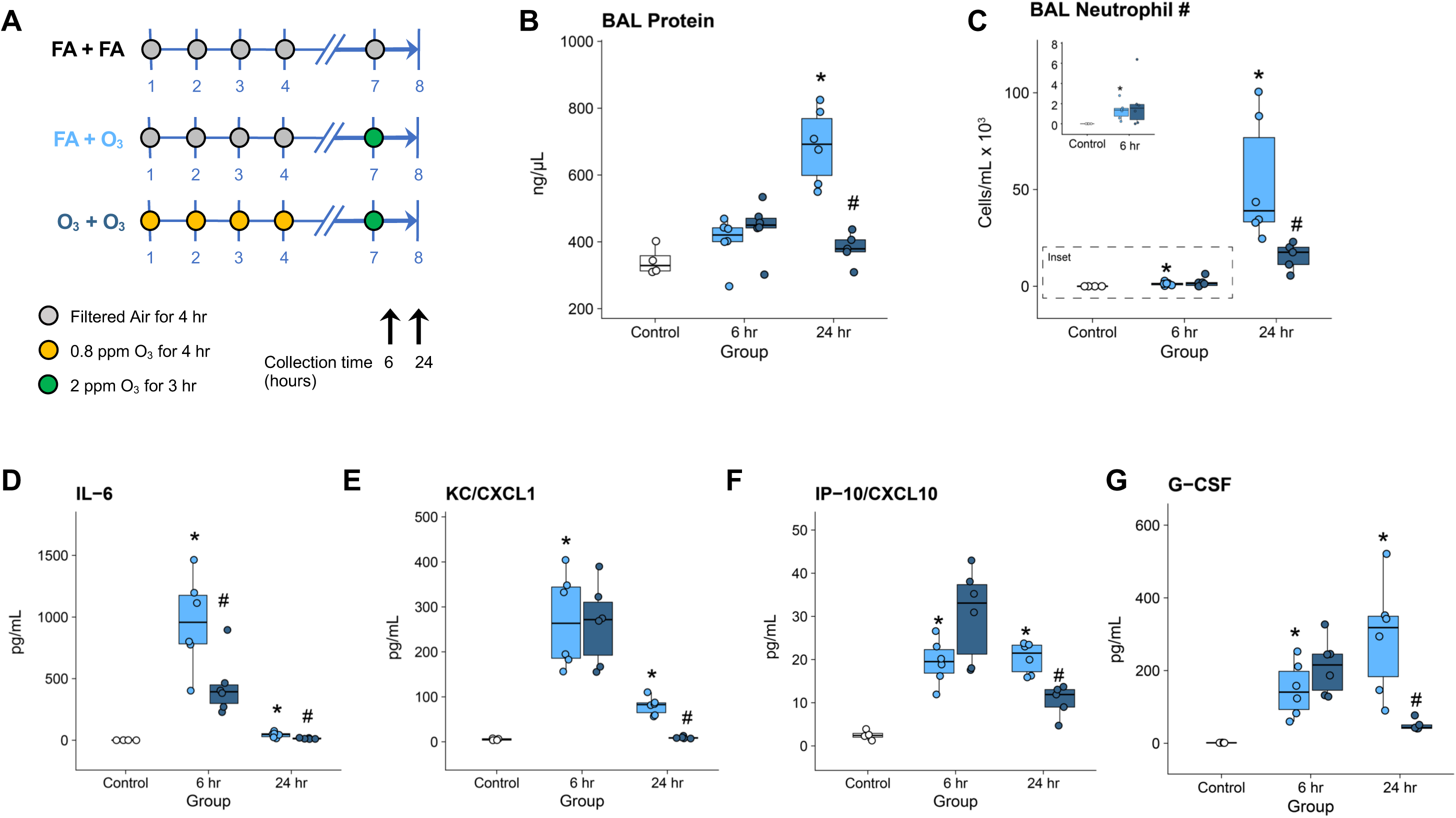
A new model of ozone tolerance in mice. **A.** Female C57BL/6NJ mice were exposed to ozone as depicted, then sacrificed 6 or 24 hours following 2 ppm challenge. **B-C**. Decreased injury and inflammation after ozone challenge in ozone pre-exposed mice as reflected by total protein (B) and neutrophils (C, D) in BAL fluid, respectively. Control mice were harvested at 24 hours. **D-G**. Altered cytokine/chemokine concentrations in BAL fluid were also observed in the ozone pre-exposed group. N=4-6/group. *p<0.05 for one-sided contrast between FA+O_3_ group vs. FA+FA group; ^#^ p<0.05 for one-sided contrast between O_3_+O_3_ vs. FA+O_3_ group.

### Lung phenotyping

At designated time points, mice were anesthetized (2 g/kg urethane) and sacrificed by inferior vena cava/abdominal aorta exsanguination. Bronchoalveolar lavage (BAL) was performed by instilling the lungs with phosphate-buffered saline containing a protease inhibitor cocktail (Roche Cat. No. 11836170001) two times, first with 0.5 mL and then with 1.0 mL. The BAL fluid was centrifuged for 10 minutes at 400 x g. Supernatant from the first fraction was saved and stored at -80^○^C for subsequent analysis of total protein and cytokines. Pellets from both BAL fractions were pooled then treated with red blood cell lysis buffer and centrifuged once more at 400 x g for 10 minutes. Pellets were resuspended in 500 µL of HBSS, and then used for total and differential cell counts.

Total BAL protein was quantified with the Qubit Total Protein Quantification kit and the Qubit 2.0 fluorometer (Thermo Scientific). BAL cytokines including IL-6, IP-10/CXCL10, KC/CXCL1, and MIP-1β, were measured using a MILLIPLEX protein immunoassay (Millipore) that was read on a Bio-Plex 200 multiplex suspension array system (Bio-Rad).

### Mucin immunoblotting

We measured the levels of soluble and insoluble MUC5AC and MUC5B protein levels in BAL fluid. BAL was performed with 0.5 mL PBS with protease inhibitors, then was spun at 4,000 rpm for 10 min at 4^○^C. The lavage was then separated into a supernatant fraction and pellet fraction, corresponding to soluble and insoluble mucin fractions, respectively. The pellet fraction was resuspended in 100 µL PBS with protease inhibitors. Western blotting for MUC5AC and MUC5B was performed using methods previously described^51,52^ with 40 µL of supernatant and pellet BAL samples using polyclonal antibodies raised against a mouse MUC5AC or MUC5B overnight at 4^○^C. Immunoblots were imaged on an Odyssey CLx imaging instrument (LI-COR Biosciences). MUC5AC was not detectable in any samples tested, therefore results are not shown for this protein.

### Antioxidant assay

We measured total antioxidant activity in lung tissue homogenates using an Antioxidant Assay Kit according to the manufacturer’s instructions (Item No. 709001; Cayman Chemical). The assay compares the ability of tissue samples to inhibit the oxidation of 2,2’-Azino-di-3-ethylbenzthiazoline sulphonate to Trolox, a tocopherol analogue. Data are reported in Trolox equivalents based on a standard curve generated from dilutions of Trolox. Previously flash-frozen (and stored at -80C) right upper lung lobes were placed in 600uL of Assay Buffer in 2mL Lysing Matrix D tubes (MP Biomedicals) and homogenized using a FastPrep-24 (MP Biomedicals). Lung tissue homogenates were centrifuged at 10,000 x g for 15 mins at 4C and 10 µl of sample was loaded in duplicate in the 96 well assay plate.

### Body temp measurements

A digital thermocouple thermometer (Model BAT-12; Physitemp Instruments) and a rectal probe were used to conduct rectal thermometry of mice pre- and post-exposure, resulting in two core body temperature measurements. The change in temperature was calculated as post-exposure minus pre-exposure temperature. Temperature data were recorded within 10 seconds (upon stabilization of the reading) and all mice were measured within 5 mins of each other.

### Chlodronate treatment

PBS or clodronate (5 mg/ml) containing liposomes (Liposoma, Amsterdam, The Netherlands) were administered to mice by oropharyngeal aspiration on day 5 of the ozone exposure protocol. The total volume administered was 50 μL, for a total dose of 250 μg clodronate. In pilot experiments, we determined that this protocol produced ∼70% reduction of BAL alveolar macrophages harvested 48 hours after clodronate treatment with minimal neutrophilia (2.3% neutrophils in vehicle treated vs. 2.9% neutrophils in clodronate treated mice, p=0.81).

### Preparation and analysis of epithelial populations by multi-color flow cytometry

Lungs were digested by intratracheal instillation via a 20-gauge catheter of 1 mL of 5 mg/mL collagenase I (Worthington Biochemical Corp, Lakewood, NJ) and 0.25 mg/mL DNase I (Sigma) prepared in RPMI media (Life Technologies, Carlsbad, CA) prior to instilling 0.5 mL of 1% (wt/vol) low melting agarose (Amresco, Solon, OH), similar to previous protocols^53–55^.

Lungs were incubated at 37°C for 30 minutes and then minced and triturated through an 18-gauge needle. Cell suspensions were then filtered through a 50 mL conical 100 μM filter (ThermoFisher, Pittsburgh, PA) before RBC lysis and stained as previously described^55^. Single cells were suspended in PBS buffer supplemented with 1 % (w/v) bovine serum albumin (Sigma) and 2 mM EDTA (Sigma), and the total cell count was determined by hemocytometer measurement. Cells (1.5 x 10^6^ cells) first underwent Fc receptor blockade with rat anti-mouse FcγRIII/II receptor (CD16/32; BD Biosciences). After blocking for 5 minutes on ice, cells were surface stained using antibodies listed in Supplemental Table 2. Cells were also concurrently stained with Zombie NIR^TM^ for live cell/dead cell discrimination. Fixed and permeabilized single-cell suspensions were subsequently stained with intracellular antibodies for Ki67 (Supplemental Table 1) to characterize differences in proliferation.

Flow cytometry was performed using a Cytoflex flow cytometer (Beckman Coulter, Brea, CA) and analyzed using CytExpert (Beckman Coulter) software, version 2.4, and our gating strategy is shown in Supplementary Figure 1. To determine the total number of a specific population in the lung, we first calculated the population’s percentage to the total live single-cell population. Next, we multiplied this percentage by the total cell count as determined by hemocytometer measurements to calculate the specific population’s total number per mouse lung as similarly described^53,54,56^.

### Preparation and analysis of leukocyte populations by multi-color flow cytometry

Murine lung leukocytes were isolated and analyzed by flow cytometry as previously described^57^. Briefly, harvested lungs were minced and digested with Liberase TM (100 µg/mL; Roche, Indianapolis, IN), collagenase XI (250 µg/ml), hyaluronidase 1a (1 mg/ml), and DNase I (200 µg/ml; Sigma) for 1 h at 37°C. The digested tissue was passed through a 70 µm nylon strainer to obtain a single cell suspension. Red blood cells were lysed with 0.15 M ammonium chloride and 1 mM potassium bicarbonate. For antibody staining of surface antigens, cells were incubated with anti-mouse FcγRIII/II receptor (CD16/CD32) for 5 min to block Fc receptors, followed by incubation with fluorochrome- or biotin-conjugated antibodies listed in Supplemental Table 1 for 30 minutes on ice. Staining with biotinylated antibodies was followed by incubation with fluorochrome-conjugated streptavidin for 20 minutes on ice. Cells were also concurrently stained with Zombie Aqua^TM^ for live cell/dead cell discrimination.

Flow cytometry data was acquired with a four-laser LSRII (BD Biosciences) and analyzed using FlowJo (Treestar, Ashland, OR) software. Only single cells were analyzed and leukocyte populations were identified using a manual gating strategy depicted in Supplementary Figure 2. In some experiments, alveolar macrophages were sorted from lung single-cell suspensions to a purity of >95% of viable cells using a four-laser FACSAria III cell sorter (BD Biosciences).

### Gene expression analysis by qRT-PCR

Total RNA was isolated from whole, right middle lung lobe tissue by column purification (RNeasy Mini Kit, Qiagen) and quantified using a Nanodrop spectrophotometer (ThermoFisher Scientific). cDNA was generated using 500 ng of total RNA (High-Capacity RNA-to-cDNA Kit, ThermoFisher Scientific). qRT-PCR reactions consisted of 5 ng cDNA sample, primer pairs (see Supplementary Table 2), and iTaq Universal SYBR Green Supermix (BioRad) and were run in triplicate using the manufacturer’s suggested cycling parameters on a BioRad CFX 384 Touch Real-Time PCR detection system. Gene expression data (Cq) were normalized to the expression of Rps20 to calculate dCq and expressed as relative quantitative values (RQV = 2^-dCq[treated]^ / 2^-dCq^ ^[control mean]^). We removed one sample from our analyses because its RNA quality was low and its Cq values were >3 SD from the mean.

### Statistical analyses

For the analyses of phenotypes in the model of tolerance (Figure 1), we defined tolerance as a statistically significant decrease in acute ozone-induced inflammation and injury as a function of ozone pre-exposure. Thus, a priori, we posited one-sided t-tests in comparisons between these two groups (i.e., O_3_+O_3_ < FA+O_3_ groups). Likewise, to examine whether ozone itself caused injury and inflammation vs. filtered air controls, we performed one-sided tests (FA+O_3_ > FA+FA groups in Figure 1A). In experiments in which the effect of macrophage depletion on tolerance was evaluated (Figure 7), we used a similar approach and a priori specified one-sided t-tests between the O_3_+O_3_ group treated with chlodronate-containing liposomes vs. O_3_+O_3_ group treated with empty liposomes (i.e., O_3_+O_3_ + chlodronate < O_3_+O_3_ + empty liposome). For most other analyses, we used standard two-sided t-tests. Some phenotype data was log-normally distributed. In these cases, we log transformed the data prior to statistical modeling. In the analyses of qRT-PCR data shown in Figure 3 and Supplementary Figure 4, we used ANOVA to test for significant differences between exposure groups followed by pairwise comparisons using Tukey’s honestly significant difference test with alpha=0.05.

### Gene expression analysis by RNA sequencing

Total RNA was extracted from two batches of flow-sorted AMs using QIAGEN RNAeasy kits per the manufacturer’s instructions. RNA integrity was analyzed using an Agilent Bioanalyzer. RIN values ranged from 8.9-9.7, indicating intact RNA. PolyA+ RNA libraries were prepared with the Roche Kapa mRNA stranded library preparation kit as per manufacturer’s instructions. Paired-end sequencing (50 cycles) was performed on an Illumina NovaSeq SP to a depth of >55M read pairs per sample by the UNC High-Throughput Sequencing Facility.

### RNA-seq data processing and analysis

Raw reads were trimmed and filtered of adapter contamination using cutadapt^58^, and further filtered such that at least 90% of bases had a quality score of at least 20 using fastx_toolkit v0.0.14. Reads were then aligned to the reference mouse genome (mm10) (Supplementary Table 3) and GENCODE vM25 transcript annotations using STAR v2.7.7a^59^, and transcript abundance was estimated using salmon v1.5.2^60^. Differential expression between ozone-exposed vs. filtered air groups was then detected using DESeq2 v1.26.0^61^ in R v3.6.0, using a design that corrected for flow cytometry batch dates. These batch effects were also removed from the VST-normalized expression values using limma v3.42.2^62^. All log2 fold-changes reported were shrunken using ashr 2.2-47^63^. Heatmaps of VST-normalized expression among differentially expressed genes were generated using tidyHeatmap v1.1.5^64^ and ComplexHeatmap^65^.

Gene ontology enrichments were then assessed using gprofiler2 v0.2.1^66^. We queried GO, KEGG and Reactome pathway databases. We also tested for enrichment of genes identified in a previous study by Choudhary et al.^38^ (GSE156799) that also examined ozone-induced changes in alveolar macrophages. To enable these analyses, we re-analyzed alveolar macrophage gene expression data from that study using DESeq2, including sex as a covariate. We also leveraged a list of M1 and M2 marker genes published in that study to examine whether these pathways were altered in our exposure model.

Raw and processed RNA-seq data have been deposited to the Gene Expression Omnibus (GEO) under accession GSE248291.

## RESULTS

### Development and characterization of a model of ozone tolerance in mice

Inspired by previous studies of ozone tolerance in rats^21,22,25^, we developed a new mouse model of ozone tolerance in which mice were pre-exposed to 0.8 ppm ozone for 4 days (4 hours/day) to induce tolerance, rested for 2 days, and then challenged with 2 ppm ozone for 3 hours (Figure 1A). Mice were phenotyped for injury and inflammation either 6 or 24 hours after the start of ozone challenge. We define tolerance here as a statistically significant decrease in inflammation and injury caused by 2 ppm ozone exposure as a function of 0.8 ppm ozone pre-exposure. We found that pre-exposure to 0.8 ppm ozone indeed resulted in significantly decreased injury and inflammation 24 hours after 2 ppm ozone challenge, as reflected by protein and neutrophils in BAL fluid, but not at the earlier time point of 6 hours (Figure 1B,C). Pro-inflammatory cytokines and neutrophil chemokines in BAL fluid largely reflected the same pattern (Figure 1D-G), but for IL-6, differences were apparent at the earlier time point. In addition to pro-inflammatory cytokines, the anti-inflammatory cytokine IL-10 was lower among ozone-pre-exposed mice at the 6 hour time point (Supplementary Figure 3).

We examined ozone’s effects on proximal and distal airways and found similar patterns of protection caused by ozone pre-exposure (Figure 2). The proximal airways of mice that were pre-exposed to filtered air and challenged with ozone exhibited a marked loss of epithelial cells resulting in a single layer of squamoid cells and exfoliated necrotic epithelial cells in the airway lumen (Figure 2E). Correspondingly, there were few FOXJ1+ ciliated epithelial cells remaining (Figure 2G). The airways of ozone pre-exposed mice, in contrast, were largely intact (Figure 2I), with ample FOXJ1+ ciliated cells present (Figure 2K). In the terminal bronchiole, we again observed denuded airways and exfoliated epithelial cells among mice pre-exposed to filtered air and challenged with ozone (Figure 2B), but the loss of FOXJ1+ ciliated epithelial cells was less severe than in proximal airways (Figure 2D). Among mice pre-exposed to ozone, the epithelium was not injured but exhibited signs of hyperplasia compared to filtered air controls (Figure 2J), likely indicative of remodeling in response to four days of ozone pre-exposure, and relatively preserved FOXJ1+ ciliated cells (Figure 2L). Overall, this model of tolerance produced the predicted effects of decreased injury and inflammation, with some time dependence for each end point. Hereafter in the text, we refer to mice pre-exposed to ozone as “tolerized”, mice pre-exposed to filtered air as “non-tolerized”; and mice exposed only to filtered air throughout the 8-day protocol are referred to as “unexposed controls”.

**Figure 2.**
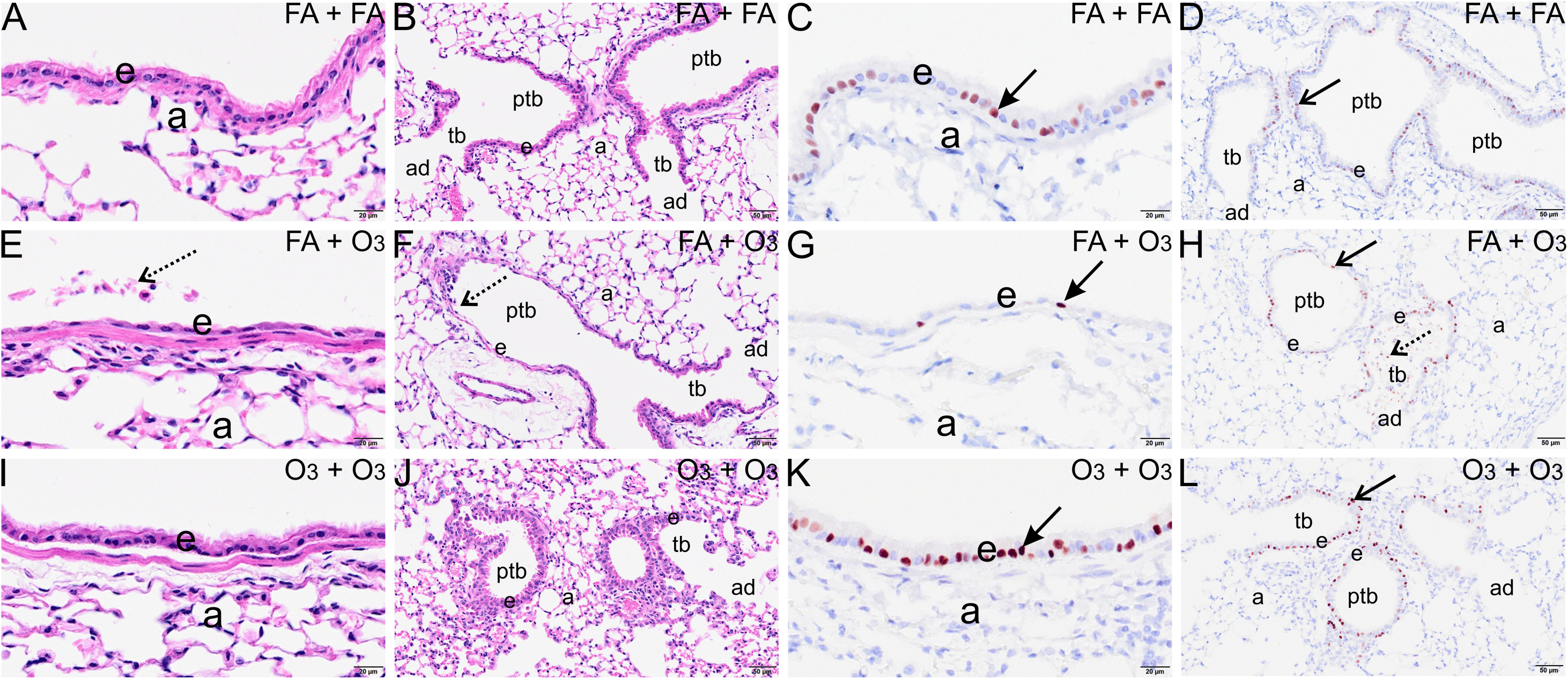
Histological analysis of large and small airways shows a protective effect of ozone pre-exposure on acute ozone-induced epithelial injury. Light photomicrographs of respiratory epithelium lining proximal large diameter axial airway (A, C, E, G, I, K) or distal small diameter pre-terminal and terminal bronchioles (B, D, F, H, J, L) in the left lung lobe of mice exposed to FA+FA (A-D), FA+O_3_ (E-H), or O_3_+O_3_ (I-L). Airway tissues were stained with hematoxylin and eosin (A, B, E, F, I, J) or immunohistochemically stained for FOXJ1 (nuclear biomarker for ciliated epithelial cells) and counterstained with hematoxylin (C, D, G, H, K, L). Stippled arrows point to exfoliated necrotic epithelial cells in the airway lumen. Solid arrows point to cells with positive nuclear staining for FOXJ1. a, alveolar parenchyma; ad: alveolar duct, e, respiratory epithelium, ptb: pre-terminal terminal bronchioles; tb: terminal bronchioles.

**Figure 3.**
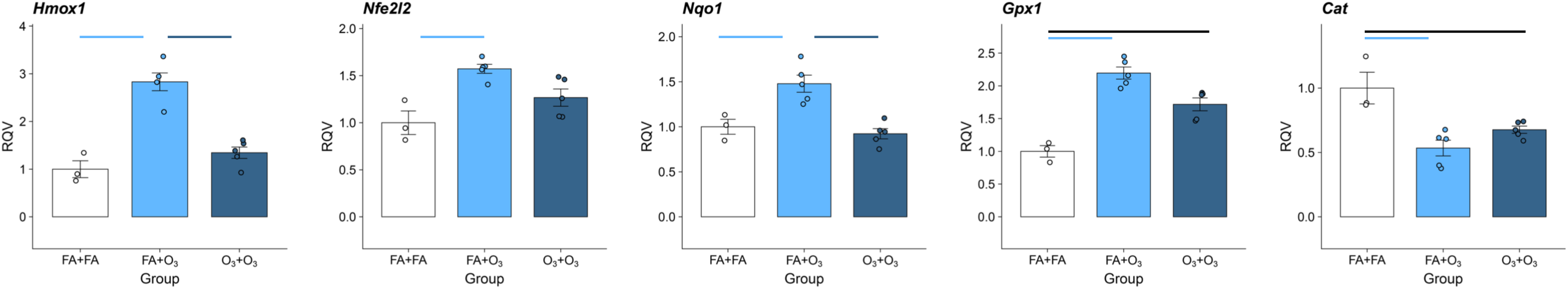
Attenuated induction of antioxidant genes among tolerized mice. qRT-PCR based measurements of antioxidant gene expression in whole lung tissue. N=4-6/group. Horizontal lines denote significantly different contrasts between indicated groups, as determined by Tukey’s post-hoc test of honestly significant differences. See Supplementary Figure 4 for analysis of other genes that showed no significant differences between groups.

### Antioxidants in the tolerance model

Given that upregulation of antioxidants has been suggested as a mechanism for tolerance, we measured the expression of genes involved in oxidative stress using qRT-PCR with RNA isolated from whole lung tissue collected 24 hours after ozone challenge (Figure 3, Supplementary Figure 4). Specifically, we evaluated the genes encoding heme oxygenase (*Hmox1*), NRF2 (*Nfe2l2)*, (NAD(P)H dehydrogenase quinone 1 (*Nqo1*), glutathione peroxidase (*Gpx1)*, Catalase (*Cat*), the genes encoding catalytic and modifier subunits of glutamate-cysteine ligase (*Gclc* and *Gclm*, respectively), and two superoxide dismutases (*Sod1* and *Sod2*). *Gclc, Gclm, Sod1* and *Sod2,* were not significantly altered in this model (Supplementary Figure 4).

In contrast, *Hmox1*, *Nfe2l2*, *Nqo1*, and *Gpx1* were significantly increased by ozone challenge in non-tolerized mice. Tolerized mice challenged with ozone exhibited significant reductions in expression in *Hmox1* and *Nqo1* compared to non-tolerized mice, and the expression of *Hmox1, Nfe2l2,* and *Nqo1* among tolerized mice challenged with ozone was not significantly different from unexposed controls. Tolerized mice also did not upregulate *Gpx1* to the same degree as non-tolerized mice, though the difference was not statistically significant, while its expression among tolerized mice was significantly elevated compared to unexposed controls. In contrast, both tolerized and non-tolerized mice downregulated *Cat* expression significantly, compared to unexposed controls, and were not significantly different from each other. Thus, by and large, these results suggest that tolerized mice do not need to mount as strong an antioxidant response compared to mice pre-exposed to filtered air, and that the degree of antioxidant gene upregulation is commensurate with injury and inflammation.

We also considered the possibility that antioxidants were upregulated prior to ozone challenge and that this conferred a degree of protection. Thus, we measured antioxidants at the time point corresponding to just before 2 ppm ozone challenge (day 7, Figure 4A). Using an assay to quantify total antioxidant capacity of homogenized lung tissue, we found that non-tolerized and tolerized mice groups did not differ (Figure 4A). We also measured antioxidant gene expression in whole lung tissue and found that most of these genes showed little or modest increases among tolerized mice, but these changes were not significant except for *Hmox1* (Supplementary Figure 5). Likewise, CCSP (*Scgb1a1*), which has putative antioxidant properties, was also not increased at the mRNA level in whole lung tissue (Figure 4C). Because mucus contains antioxidants, increased mucus production has been posited as a potential protective mechanism against oxidative stress^29,30^. Thus, as a complementary approach to measuring antioxidants, we measured the quantities of both the soluble and insoluble forms of the major secreted airway mucins, MUC5AC and MUC5B, in BAL fluid. Both soluble and insoluble forms of MUC5AC were undetectable (not shown). MUC5B was detectable, but neither soluble nor insoluble levels of MUC5B were different between non-tolerized and tolerized mice (Figure 4D,E). In total, these data argue against upregulation of antioxidants, CCSP, and mucus as protective mechanisms in this model of tolerance.

**Figure 4.**
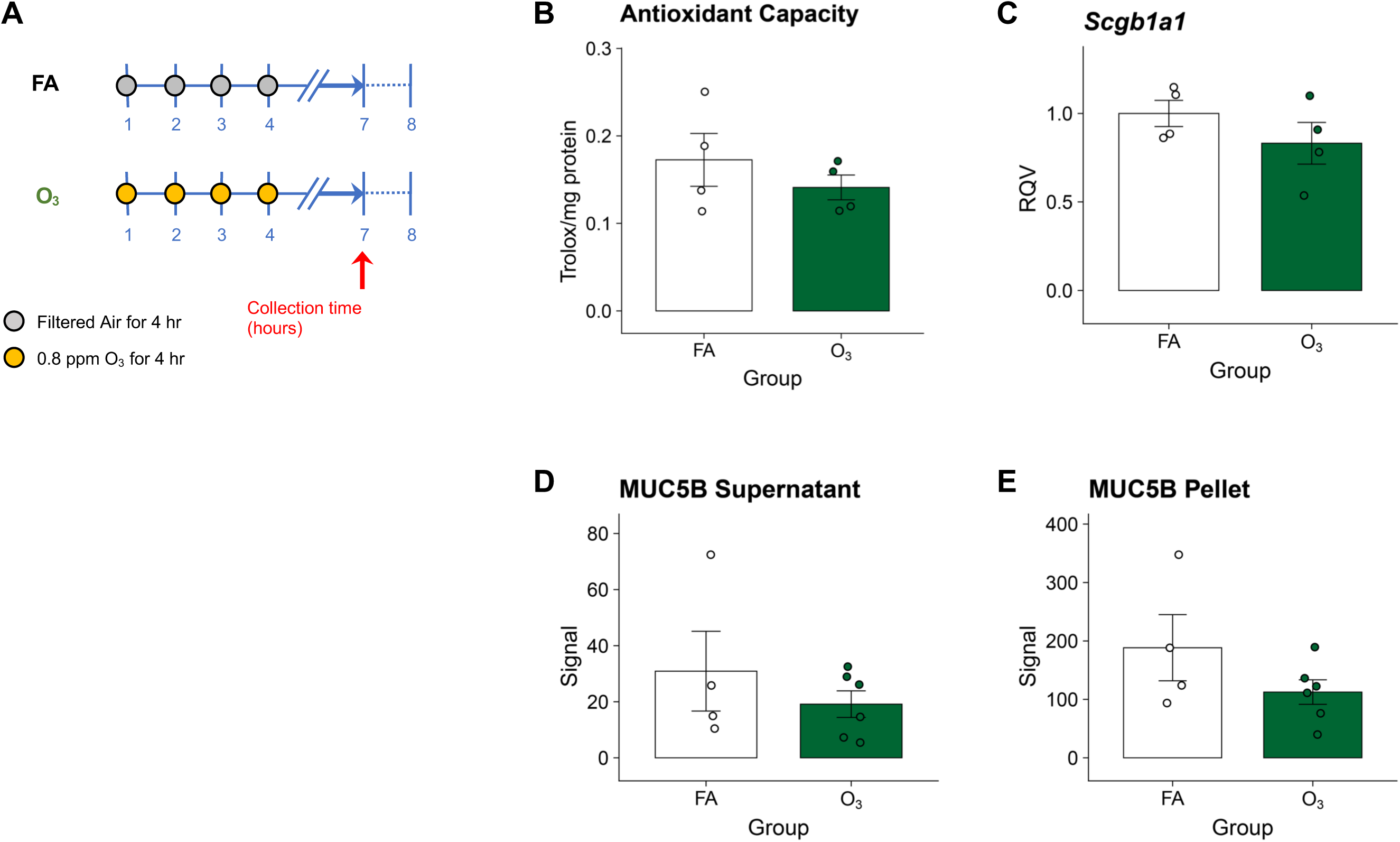
Ozone tolerized mice do not upregulate antioxidants, CCSP, or mucin proteins. **A.** Exposure paradigm with mice phenotyped at the timepoint corresponding to just prior to ozone challenge shown in Figure 1. **B**. Total antioxidant capacity of lung tissue samples, normalized to total protein. **C.** CCSP (*Scgb1a1*) gene expression in whole lung tissue. **D** and **E.** MUC5B protein levels in supernatant (D) and pellet fractions (E) of BAL. MUC5AC was not detectable. N=4-6/group.

### Evaluation of ozone uptake in the tolerance model

We asked whether tolerance could be explained by differences in ozone uptake. It is well known that during ozone exposure, mice decrease their ventilation, ostensibly to limit inhalation of the toxic gas^67–69^. As a consequence of decreased ventilation, metabolism is decreased and core body temperature (T_co_) decreases. Slade and colleagues have shown that mice that exhibit greater decreases in T_co_ ultimately take on a lower dose^68^. Thus, T_co_ is a surrogate of ozone uptake and dose. Additionally, decreasing T_co_ may also be protective by virtue of slowing enzymatic reactions that produce toxic intermediates or reaction byproducts^70^. In our model, we found that the first exposure to ozone caused a marked decrease in body temperature, roughly 10 degrees C (Figure 5), consistent with prior studies^68,71^. With each additional exposure, this effect diminished such that after the fourth exposure, non-tolerized and tolerized mice were not different in T_co_ following exposure. Following two days of rest and challenge with 2 ppm ozone, both non-tolerized and tolerized mice had significantly different decreases in T_co_, but the effect was quite modest in size, with the two groups differing by only 1.1 degrees C. Additionally, the T_co_ decrease was less pronounced in tolerized mice. These results argue against a major role for thermoregulation, and thus altered ozone uptake and/or production of reaction byproducts, in the blunted inflammation and injury responses caused by 2 ppm ozone challenge.

**Figure 5.**
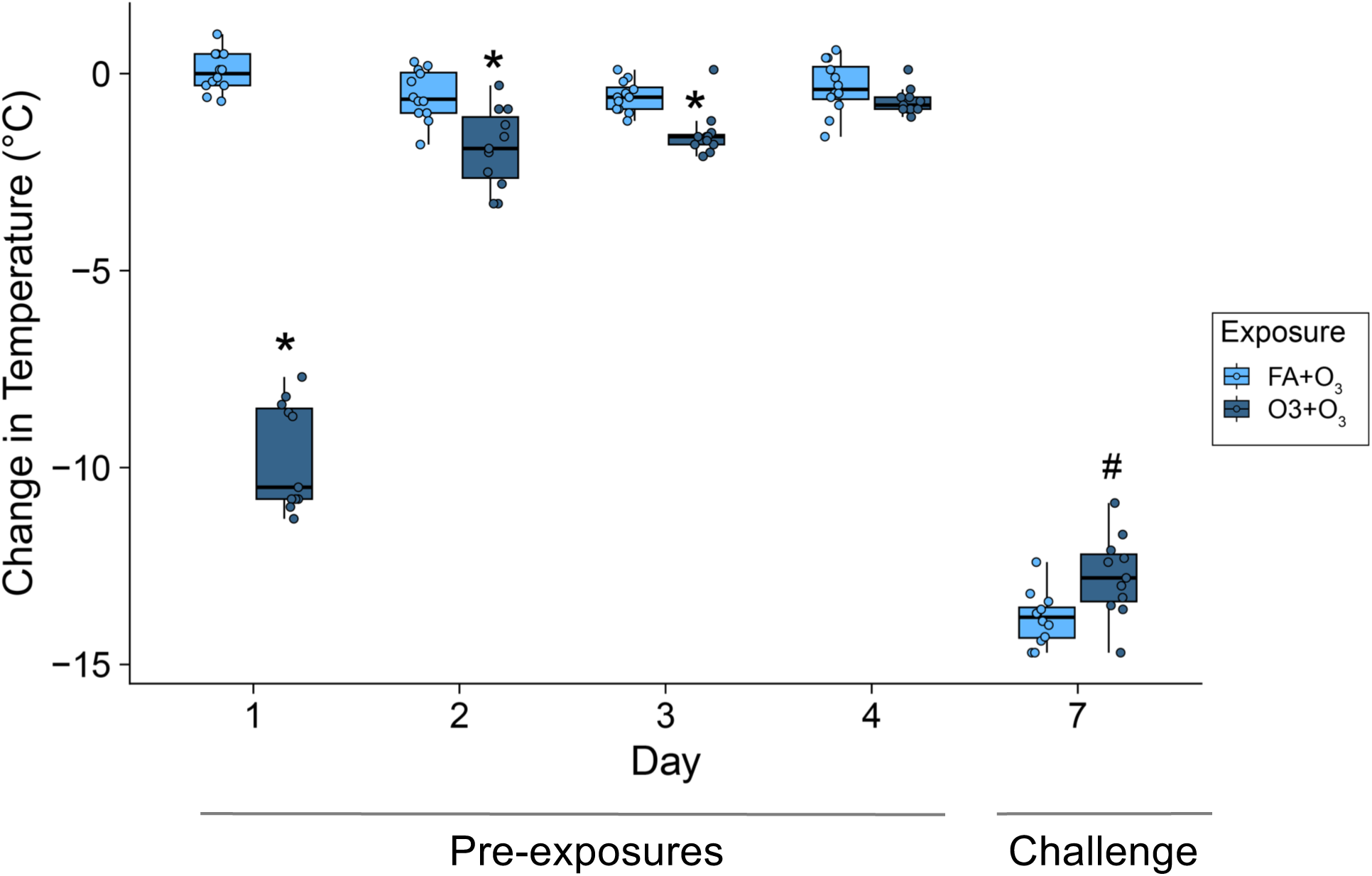
Repeated ozone exposure causes adaptive changes in body temperature but this effect largely erodes prior to acute ozone challenge. Mice were exposed to ozone as shown in Figure 1 with body temperature recorded immediately prior to and immediately after each exposure. Data represent the change in body temperature between these two measurements. N=11-12/group. * p<0.001 vs. FA+O_3_ group; ^#^ p<0.01 vs. FA+O_3_ group.

### Epithelial cell populations in tolerized vs. non-tolerized mice prior to ozone challenge

Using flow cytometry, we evaluated whether tolerized vs. non-tolerized mice exhibited differences in epithelial cell populations in the lung prior to ozone challenge (day 7). First, we measured the frequencies of epithelial cell populations, including club cells, ciliated cells, alveolar type 2 (AT2) cells, and AT2 cells transitioning to an AT1 phenotype (AT2→AT1). Total epithelial cell numbers (CD326+ cells) were not significantly different between non-tolerized and tolerized mice (Supplementary Figure 6A). The numbers of ciliated cells, club cells and AT2→AT1 cells were also not different between groups, but AT2 cell number was lower on average among tolerized mice (Supplementary Figure 6B). When expressed as a percentage of all epithelial cells, AT2 frequency was significantly decreased (Figure 5A). While AT2 frequency was lower, these cells exhibited a higher proliferative index (Ki67+) (Figure 5B), consistent with prior studies^21,23^. The frequency of AT2→AT1 cells was also significantly higher in tolerized mice (Figure 5A). Surprisingly, the frequency of ciliated cells was increased among tolerized mice (Figure 5A), though we did not observe an increased proliferative index among this population of airway epithelial cells.

### Evaluation of the role of alveolar macrophages in ozone tolerance

We then examined the potential role of AMs in ozone tolerance. First, we measured the frequencies of AMs in tolerized vs. non-tolerized mice prior to ozone challenge (day 7) and found that they were not different, whereas we did detect small but statistically significant differences in the frequencies of other leukocytes, including interstitial macrophages, T-cells, and a subset of dendritic cells (Supplementary Figure 7). All other leukocyte populations were not different in frequency between groups.

To functionally assess the contribution of AMs to tolerance, we utilized the same ozone exposure paradigm as earlier described, but 48 hours prior to 2 ppm ozone challenge, we administered liposomes containing clodronate via the airways to deplete AMs (Figure 6A). Based on prior studies^43,44^, we predicted that depletion of AMs using clodronate would result in loss of tolerance. As such, we tested for increased injury and inflammation in clodronate treated mice.

**Figure 6.**
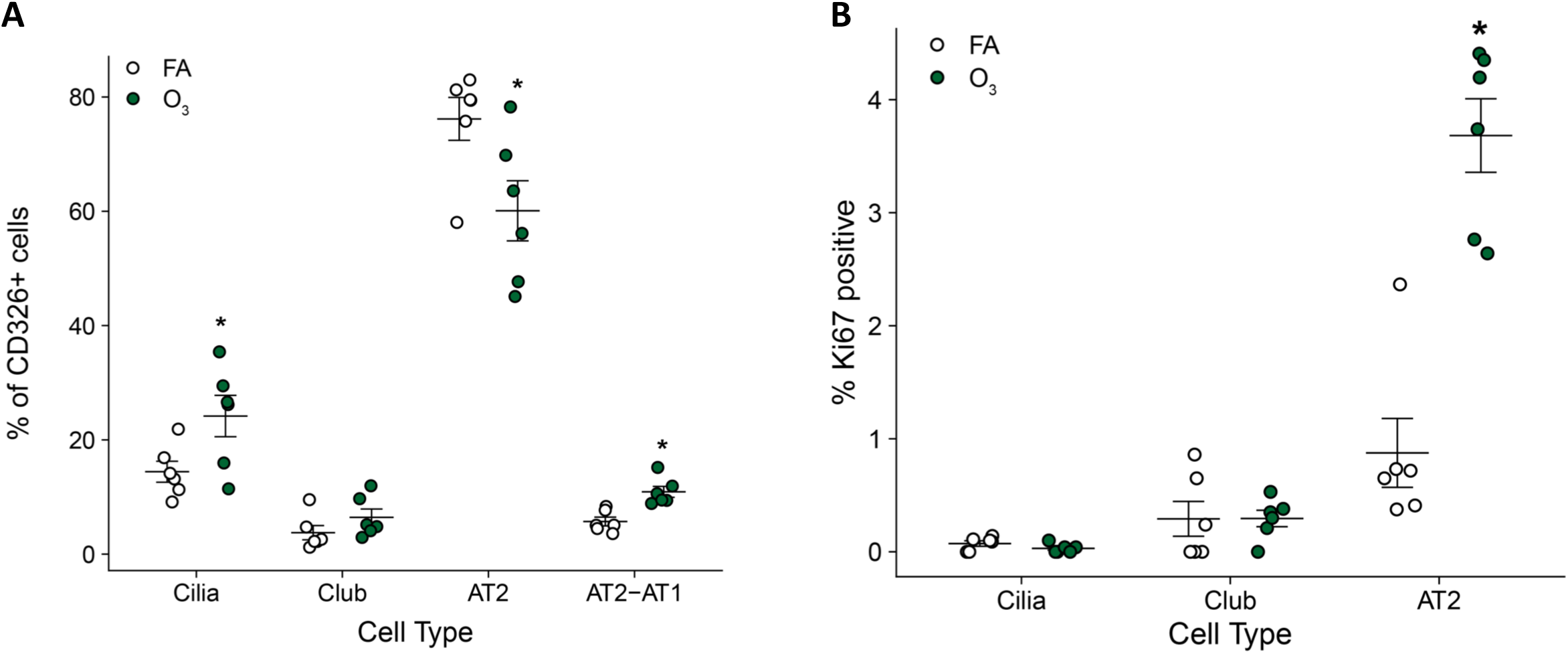
Changes in epithelial cell type frequencies and proliferation in ozone tolerized mice. Flow cytometry-based determination of epithelial types in whole lung digests. Mice were exposed to 0.8 ppm ozone for four days and harvested on day 7 as depicted in Figure 5. **A.** Frequency of epithelial sub-types relative to all epithelial cells (CD326+). **B.** Ki67 expression in club cells, ciliated cells, and AT2 cells. N=5-6/group. * p<0.05 vs. FA control group. Data for Ki67 expression in AT2>AT1 cells is not shown because they are extremely low in frequency.

**Figure 7.**
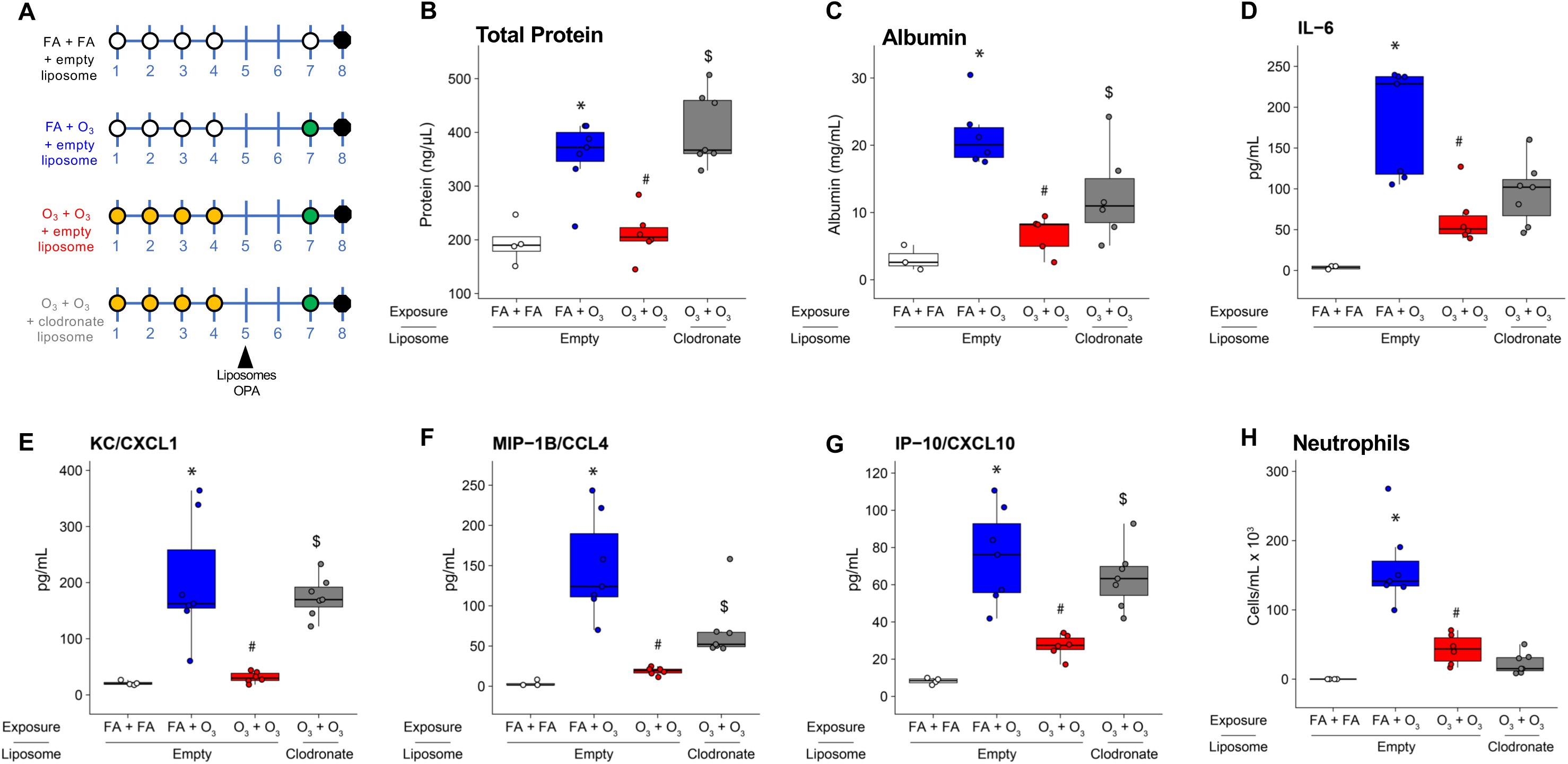
Depletion of AMs using clodronate diminishes ozone tolerance phenotypes. **A.** Experimental paradigm in which female C57BL/6NJ mice were exposed to ozone or filtered air as depicted and treated with liposomes (empty vs. clodronate containing) via oro-pharyngeal aspiration. **B-C.** Clodronate treatment caused a return of ozone-induced injury as reflected by total protein (B) and albumin (C) in BAL fluid. **D-G**. Clodronate-induced changes in BAL IL-6 (D), CXCL1/KC (E), CCL4/MIP-1b (F), and CXCL10/IP-10 (G). **H.** Clodronate did not affect recruitment of neutrophils to the airspace. N=4-7/group. Statistical tests among empty liposome treated mice: *p<0.05 for one-sided contrast between FA+O_3_ group vs. FA+FA group; ^#^ p<0.05 for one-sided contrast between O_3_+O_3_ vs. FA+O_3_ group. $ p<0.05 for one-sided contrast between O_3_+O_3_ group treated with chlodronate-containing liposomes vs. O_3_+O_3_ group treated with empty liposomes.

**Figure 8.**
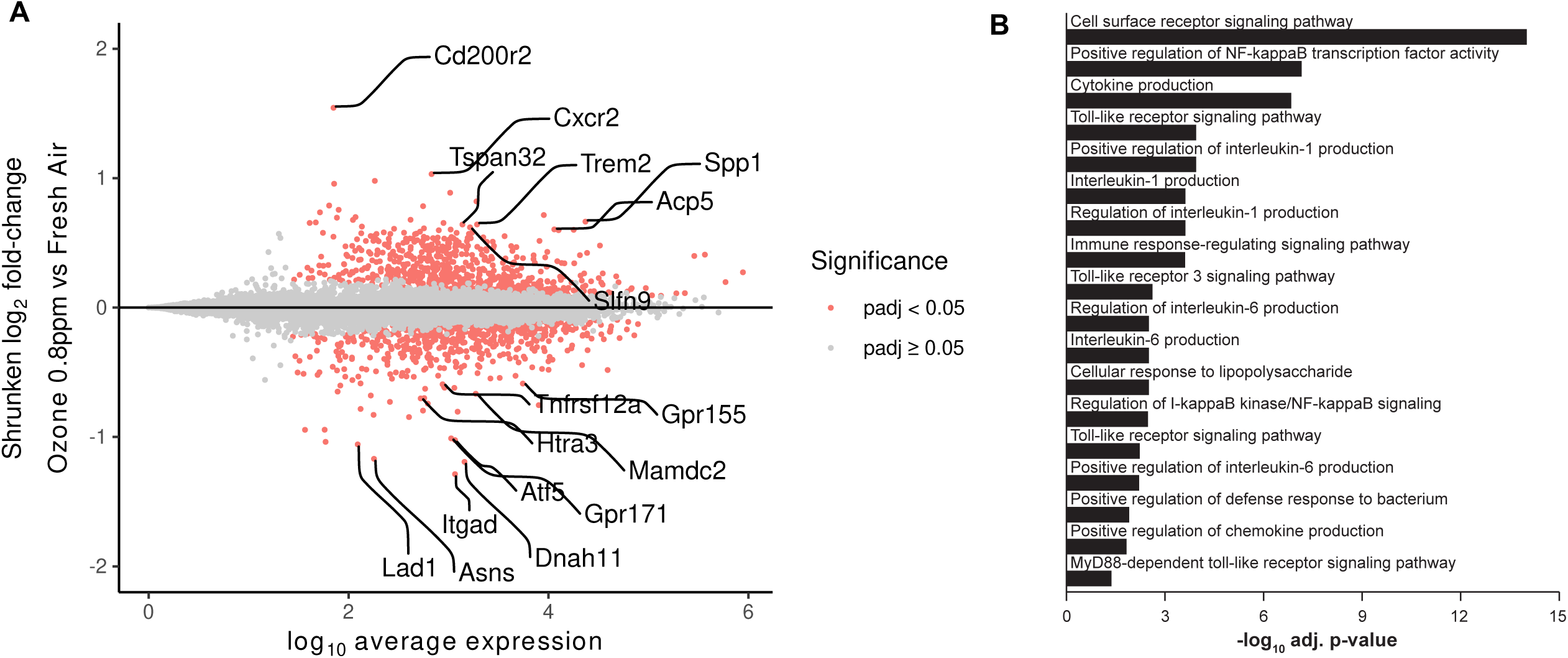
Ozone tolerance causes a shift in AM gene expression toward decreased responsiveness. **A.** MA plot showing differential gene expression as a function of average gene expression (x-axis) and log2 fold change (y-axis). DEGs with FDR<1x10^-10^ and log2 fold change > |0.58| (corresponding to fold change >|1.5| are labeled. N=6/group. **B.** Select pathway enrichment results of down-regulated DEGs (full results of pathway analysis are given in Supplementary Tables 5-6).

Indeed, compared to tolerized mice exposed to empty liposomes (vehicle control), clodronate-treated mice exhibited increased injury as reflected by total protein and albumin in BAL (Figure 6B, C). Likewise, pro-inflammatory cytokines and chemokines including IL-6, IP-10/CXCL10, KC/CXCL1, and MIP-1β, were all significantly increased compared to vehicle controls (Figure 6D-G). Despite the increase in neutrophil chemokines, however, we did not detect an increase in BAL neutrophils (Figure 6H).

### Characterizing the transcriptome of AMs tolerized to ozone

We then asked whether the transcriptome of AMs of tolerized mice exhibited significant changes compared to non-tolerized mice, specifically looking for gene expression differences that could help provide insight into the mechanism of tolerance. We isolated RNA from highly purified AM populations from tolerized and non-tolerized mice, then performed RNA-seq analysis of gene expression. Overall, we detected 1,479 differentially expressed genes (DEGs) at an false discovery rate (FDR) <0.05; of these, 62 exhibited fold changes >1.5 (32 upregulated and 30 downregulated) (Figure 7A and Supplementary Table 4). Perhaps unsurprisingly, the DEGs we detected here overlapped with DEGs detected in a previous study of AMs in which mice were exposed to 0.8 ppm ozone for 14 days^38^, and the vast majority of DEGs showed consistent patterns of up or down regulation (Supplementary Figure 8). Among the most significantly and strongly upregulated DEGs in our study were: *Cd200r2*, a member of a family of genes involved in AM-epithelial cross talk; *Spp1*, also known as osteopontin, which has previously been shown to be upregulated after ozone exposure in both mice^37,38,72^ and humans^73^; *Trem2*, which has been shown to suppress TLR-mediated cytokine production^74^ as well as bind lipids^75^; *Cxcr2*, a receptor for CXC chemokines; and *Acp5*, which encodes an enzyme that previously was shown to negatively regulate cytokine responses in peritoneal macrophages^76^.

The roles of the most strongly down-regulated genes in relation to alveolar macrophage function were less obvious, though *Itgad* (Cd11d) is an integrin previously shown to influence response to systemic infection^77^, and *Atf5* is a transcription factor known to be involved in establishing AM identity^78^. We turned to pathway enrichment analysis to infer potential mechanisms related to ozone tolerance (Supplementary Tables 5-6), querying Gene Ontology, KEGG, and Reactome databases. The most striking finding using the list of downregulated DEGs was enrichment of pathways related to cell signaling, including toll-like receptor (TLR) signaling, MYD88-dependent signaling, and NF-kB mediated signaling (Figure 7B). Pathways related to production of proinflammatory cytokines (IL-1 and IL-6) were also represented. Genes in these sets of pathways showed extremely consistent downregulation across biological replicates (Supplementary Figure 9). Collectively, these findings argue that repeated ozone exposures caused a down regulation of AM signaling pathways and cytokine production, resulting hypo-responsiveness to acute ozone challenge.

## DISCUSSION

We built a new model of ozone tolerance in mice that recapitulated findings from prior studies using rats, namely that pre-exposure to relatively low concentration (0.8 ppm) provides a degree of protection against injury and inflammation caused by challenge with high concentration (2 ppm) ozone. The degree of protection conferred by ozone pre-exposure was impressive and consistent across of a set of metrics of injury and inflammation. In addition to the diminished inflammation and injury responses to acute ozone challenge, we also found that tolerized mice exhibited significantly reduced induction of antioxidant genes, further indicating that responses overall were muted by ozone pre-exposure.

Having established the tolerance model’s utility, we evaluated previous hypotheses about tolerance mechanisms, including the role of antioxidants because several studies in rats have reported that during the development of adaptation or tolerance to ozone or other oxidants, antioxidants are upregulated^18,25,26,79,80^. However, the results of two studies argue against a causal role for antioxidants in conferring protection. First, Nambu et al.^25^ found that antioxidant upregulation occurred after tolerance had developed. Second, a study examining adaptation to chlorine gas in mice found that glutathione levels were not upregulated after repeated exposure and, further, *Nfe2l2* (NRF2) knockout mice exhibited equivalent degrees of adaptation as compared to wildtype mice. In our model, antioxidants and other putative protective factors, namely CCSP and mucus levels, were not upregulated prior to 2 ppm ozone challenge. Thus, our data also argue against an important role for these factors in protection against injury and inflammation in this model of tolerance.

In addition to measuring antioxidants, CCSP and mucus, we also investigated whether tolerance was associated with changes in the thermoregulatory response to ozone. Upon inhalation of ozone, mice exhibit a sensory nerve mediated decrease in ventilation and corresponding decreases in basal metabolic rate and core body temperature (T_co_), which likely limit toxicity by reducing ozone uptake and/or altering its toxicokinetic and toxicodynamic profile^68^. If a change in these physiological responses were responsible for tolerance, we would have expected a greater T_co_ drop in response to 2 ppm ozone challenge in tolerized mice. However, we observed that tolerized mice had a smaller drop in T_co_ than non-tolerized mice. The reason underlying the modest difference in T_co_ between tolerized vs. non-tolerized mice remains to be determined. One could postulate that pre-exposure to 0.8 ppm ozone caused changes in sensory nerve activation by various stimuli including ozone reaction products, inflammatory mediators, etc. However, if this was the case, one would have expected differences in injury and inflammation at early time points following 2 ppm ozone challenge because the effect of ozone on T_co_ occurs almost immediately after the start of exposure and reaches its nadir during exposure^68^. Rather, except for BAL IL-6, all other metrics of response at the early time point of 6 hours were not different between groups, whereas clear differences were present at 24 hours. Collectively, the directionally inconsistent effect of ozone pre-exposure on T_co_ responses to 2 ppm ozone challenge and lack of differences in injury and inflammation at the early time point argue against an important role for thermoregulation in tolerance.

After evaluating previous hypothesis for tolerance as well as dosimetry, we turned our attention to AMs, which have long been known to play a role in acute ozone responses^33–41^. Based on prior studies of adaptation/tolerance to other stimuli^43,44^, we hypothesized that AMs may be key players in ozone tolerance as well. Similar to studies of LPS tolerance in bone marrow derived macrophages^43,81–83^, we found that tolerance to ozone was associated with down regulation of cell signaling pathways that utilize Toll-like receptors, MYD88, and NF-kB and culminate in cytokine production. Our results also mirror those of Allard et al.^44^ who studied AMs in the context of adaptation to chlorine gas, also a strong oxidant, and found numerous changes in the AM transcriptome. Collectively, these studies show that repeated exposures to a given stimulus induce a state of hypo-responsiveness that has presumably evolved to prevent over-activation of immune response pathways. That similar findings have been found with these different exposures suggests that a potentially common feature is activation of pattern recognition receptors, most notably Toll-like receptors. For ozone and chlorine, the ligands for these receptors are likely secondary byproducts of reactions with components of the airway surface lining fluid and/or cell membrane or other danger associated molecular patterns (DAMPs). Previous studies have also linked tolerance to LPS in macrophages to epigenetic changes^43,84,85^. Thus, it would be worthwhile to assess how ozone tolerance alters the epigenome of AMs. That said, the lack of durability evidenced in previous study of ozone tolerance study^21^ argues against long-lasting reprogramming (i.e., innate immune memory).

Though pathway enrichment analysis of AM DEGs revealed a striking down regulation of genes involved in receptor mediated signaling and the downstream production of pro-inflammatory cytokines, we did find that some genes were upregulated by four days of ozone exposure. Perhaps most notably, *Spp1* (osteopontin) was upregulated, and this gene has been found to increase after a single ozone exposure in both mice^37,38,72^ and humans^73^. Thus, as is the case for LPS^43^, it appears that not all genes tolerize. Johnston et al. showed that *Spp1* knockout mice exhibit decreased injury and inflammation after acute ozone exposure^72^, thus sustained upregulation after repeated ozone exposure may represent one additional protective mechanism of tolerance. Additionally, *Tnfaip3* and *Acp5* were upregulated, both of which are known to be negative regulators of cytokine production^76,86^. The gene showing the greatest degree of upregulation in tolerized mice was *Cd200r2*. Comparatively little is known about this gene relative to its family members *Cd200r* and CD200 ligand. This receptor-ligand pair mediates immunosuppressive cross talk between epithelial cells and AMs^87^. As such, perhaps upregulation of *Cd200r2* is indicative of an attempt to suppress ozone-induced inflammatory signals emanating from epithelial cells. In total, it appears that ozone tolerance is a function of both down regulation of pro-inflammatory signaling pathways and upregulation of genes/pathways that suppress inflammation.

While depleting AMs from tolerized mice had clear effects on acute ozone-induced injury, neutrophil numbers were not likewise affected. This lack of effect is particularly intriguing given that hallmark neutrophil chemokines (CXCL1/KC and CCL4/MIP-1B) were increased by depletion of tolerized AM. Clodronate was recently shown to have unexpected effects on neutrophils as opposed to just AM^88^, offering one potential explanation. That said, our finding may also imply that epithelial cells also play a role in tolerance. In agreement with previous studies^21,23^, we found that tolerance was associated with a change in AT2 cell proliferation, which likely indicates that ozone pre-exposure caused a degree of AT1 injury and death, prompting AT2 cells to proliferate and then differentiate into AT1 cells to restore injured alveoli. How this proliferative effect confers protection is not immediately apparent. We suspect that newly formed AT1 cells may be less susceptible to ozone toxicity, but the underlying mechanisms remain to be determined. There is also prior evidence human large airway epithelial cells adapt to ozone^89^. Thus, further studies to investigate the role of epithelial cells in both the large airways and alveoli are warranted.

We focused our study on tolerance, but several previous studies have addressed adaptation to inhaled toxicants, that is, the effect of repeated exposures vs. a single exposure to the same concentration (and no challenge with a higher concentration of the toxicant). We presume that the induction of AM hypo-responsiveness plays a role in both adaptation and tolerance. Intriguingly, one such study by Gordon and colleagues^90^ found that classical inbred strains of mice differ in their propensity to adapt to repeated zinc oxide exposure. In particular, they found that BALB/cByJ adapted to zinc oxide while DBA/2J did not. Similar patterns of response were found when using 0.6 ppm ozone but not LPS. Thus, the genetic basis of adaptation to these different exposures is at least partly distinct. Based on our previous study documenting strain-dependent transcriptional responses in AMs to acute ozone in vivo^42^ and a study examining transcriptional responses in vitro of bone marrow-derived macrophages to multiple agonists^91^, we postulate that the genetic basis tolerance/adaptation may be elucidated in the future through genetic analyses of AM transcriptional programs of strains that differ in the propensity to adapt/tolerize.

Eventually, it will be important to evaluate whether adaptation to ozone in humans, which is well documented in controlled exposure studies^8–17^, is also a function of shifts in the AM transcriptome. We note also that in more than one of these studies, there were subjects who failed to adapt upon repeated ozone exposure^16,17^. Naturally, this leads to the question of whether differences in AM transcriptional responses to repeated ozone exposure is a determinant of failure to adapt. Given that the strain-dependence of adaptation in mice clearly indicates a genetic contribution^90^, one might further speculate that genetically determined variation in AM responses, as has been reported in a prior in vitro study^92^, may also be important in ozone adaptation.

In conclusion, we have developed a mouse model of tolerance and shown that alveolar macrophages play a key role in mediating the decreased responsiveness of ozone pre-exposed mice to acute ozone challenge. In addition to a key role of AMs, proliferating AT2 cells are also likely involved in conferring protection, as others have shown previously.

## Acknowledgements

We thank Dr. Dan Costa for many helpful conversations about tolerance and adaptation, Dr. Adelaide Tovar for brainstorming sessions, Daniel Vargas and Rowan Merritt for their assistance in performing mouse experiments, Alessandra Livraghi-Butrico for assistance with mucin immunoblotting experiments, the UNC Flow Cytometry Core Facility for assistance with cell sorting, and Amy Porter of the Michigan State University Investigative Histopathology Laboratory for assistance with histology. This research was funded by NIH Grant R01ES024965, K08ES029118, a UNC Center for Environmental Health and Susceptibility Pilot Project Award (P30ES010126), and a T32 training grant (ES007126-35).

## Supplementary Figure Legends

**Supplementary Figure 1.**
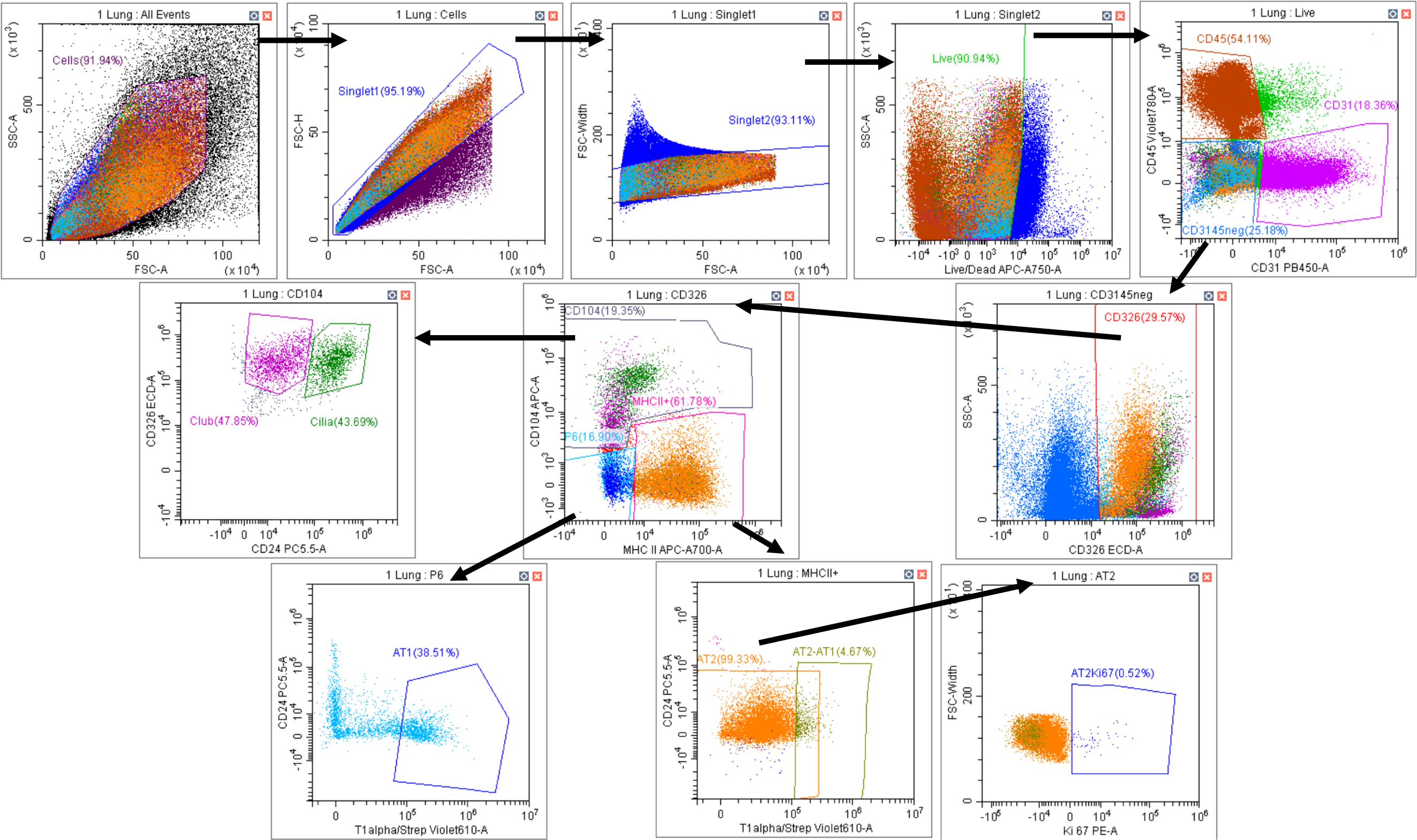
Flow cytometry gating strategy used to identify and quantify epithelial subtypes in the lung.

**Supplementary Figure 2.**
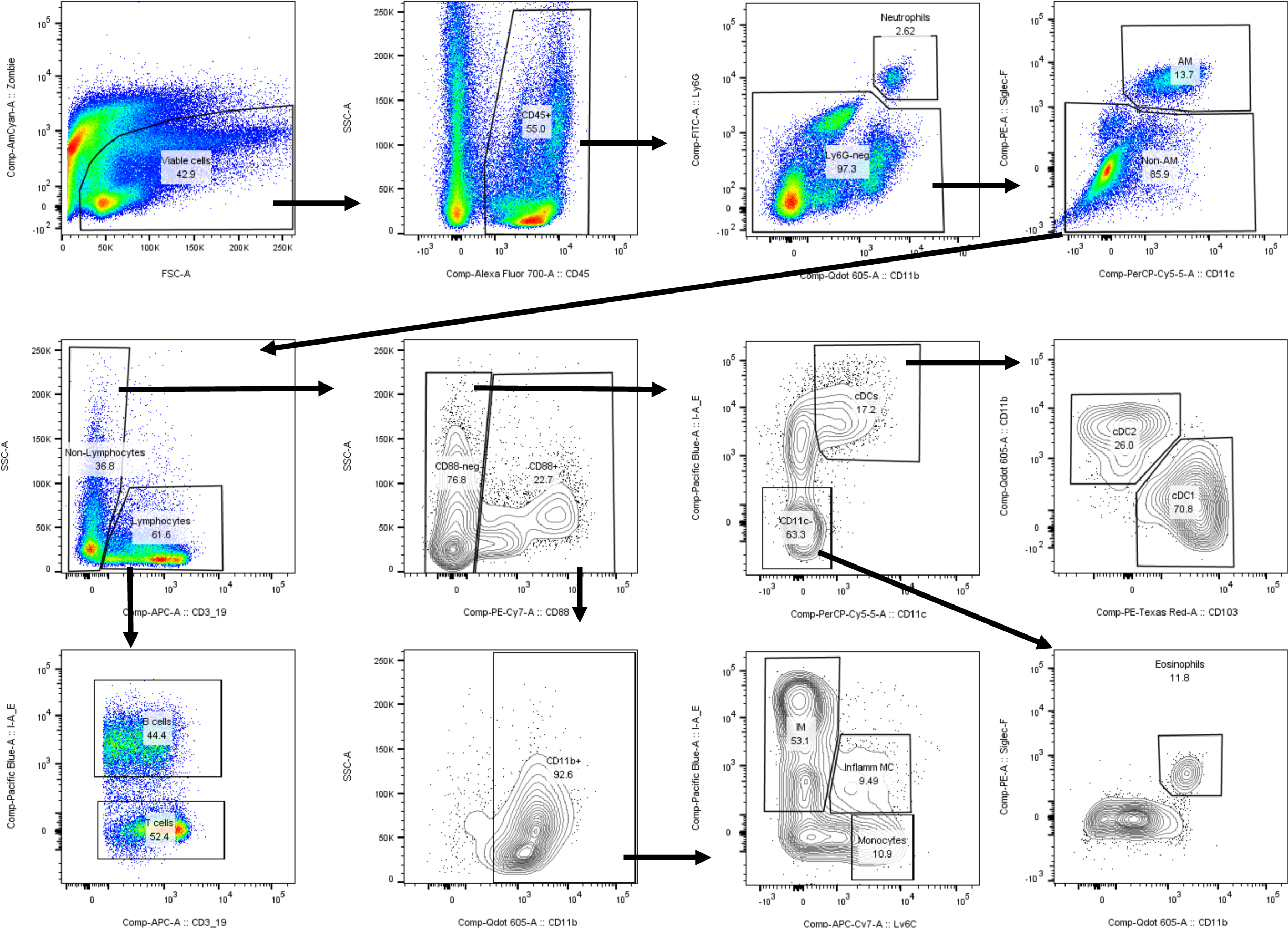
Flow cytometry gating strategy used to identify and quantify leukocyte populations in the lung. Whole lungs were digested and prepared for flow as described in the methods.

**Supplementary Figure 3.**
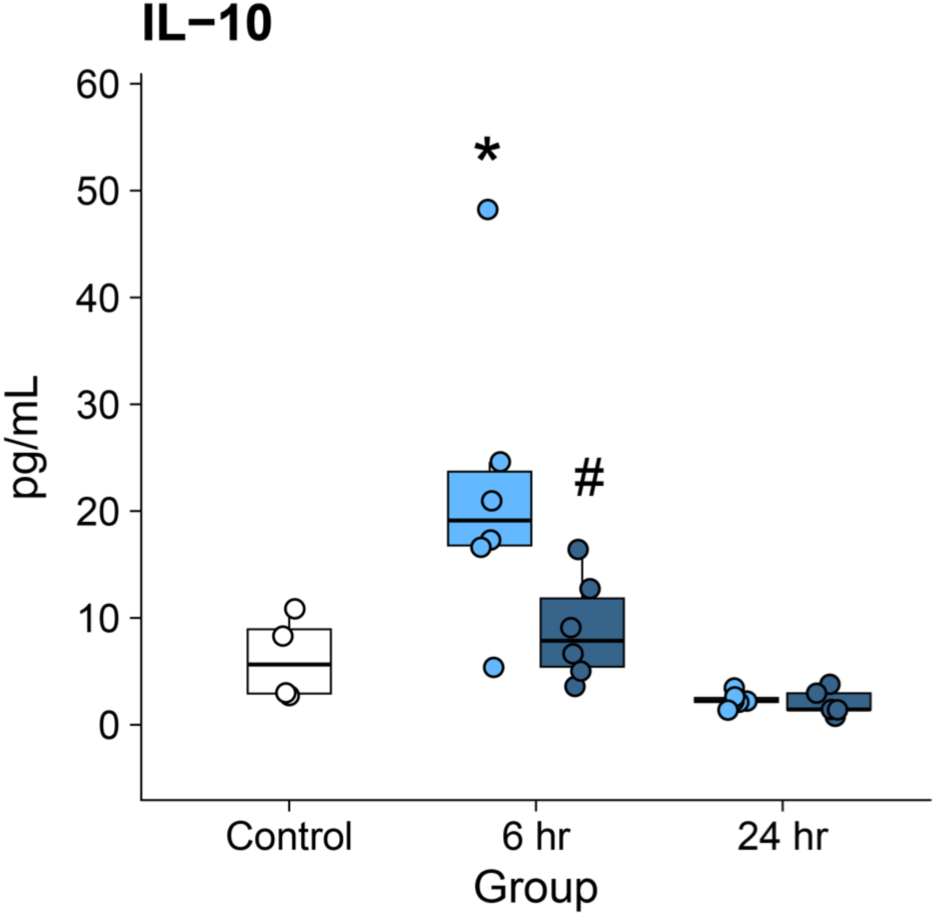
BAL IL-10 concentrations among from mice shown in Figure 1.

**Supplementary Figure 4.**
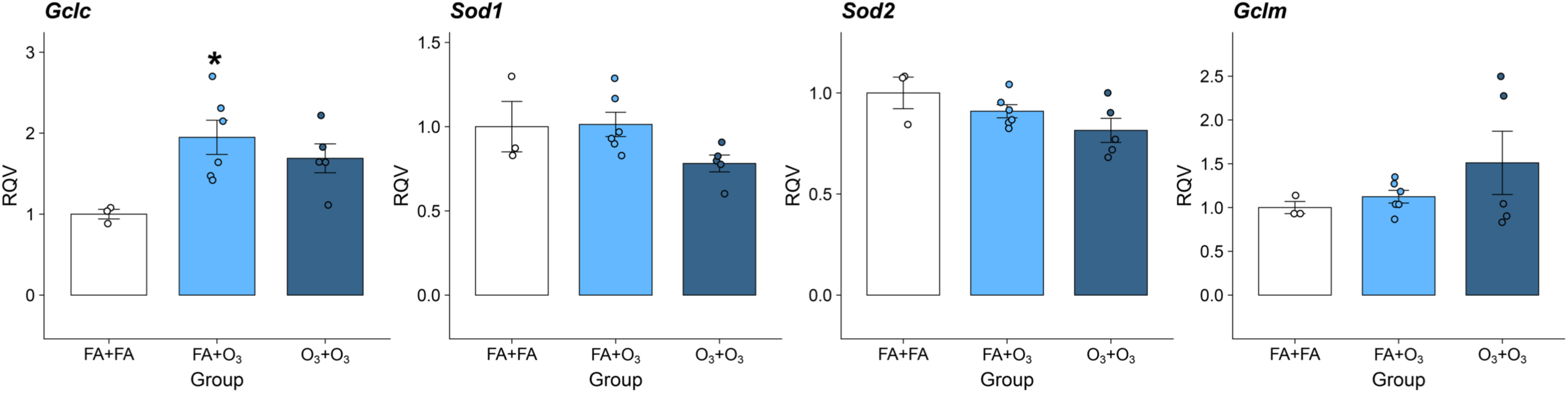
Additional qRT-PCR based gene expression measurements of antioxidant genes.

**Supplementary Figure 5.**
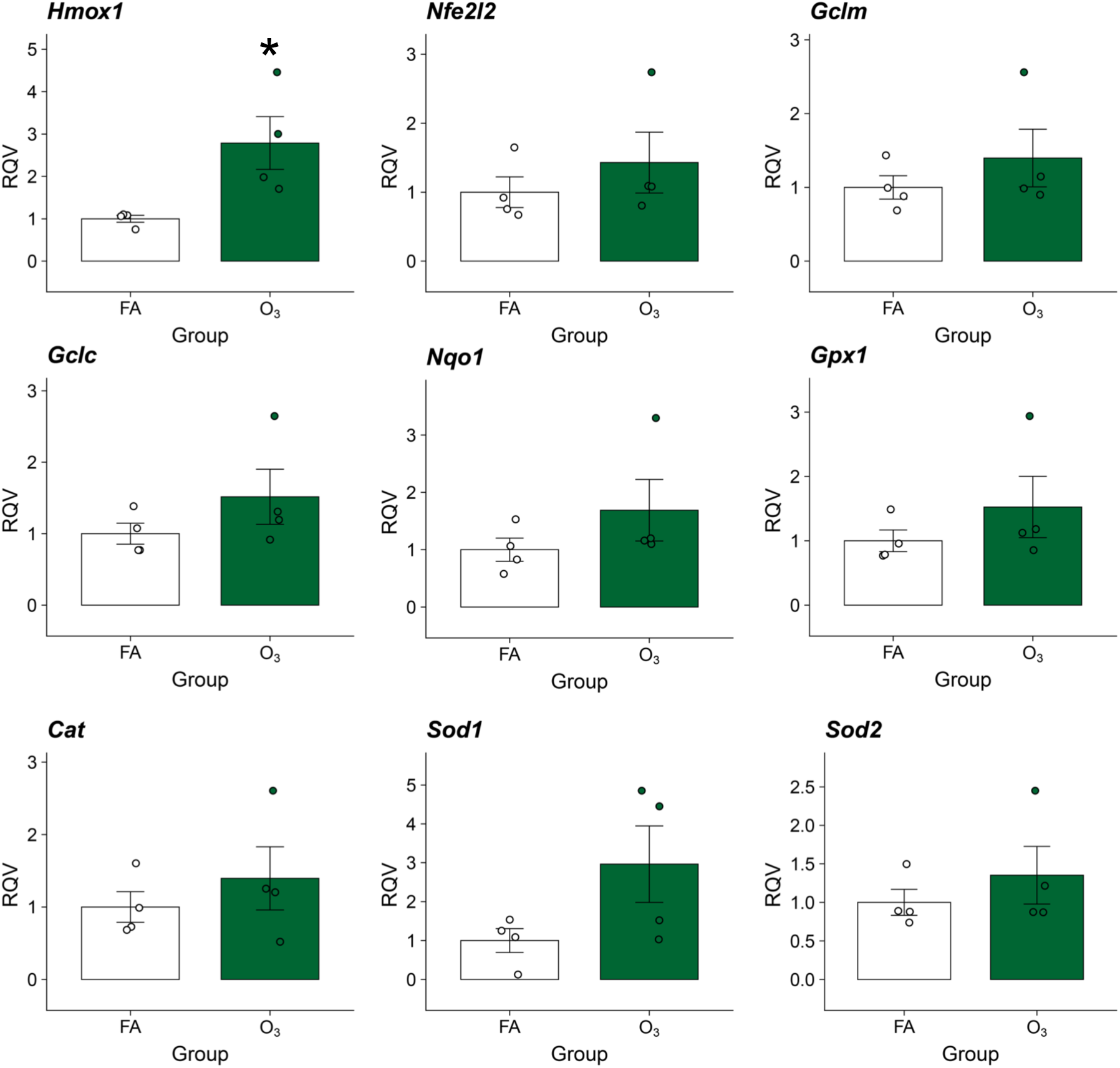
qRT-PCR based gene expression measurements of antioxidant genes in lungs from mice pre-exposed to ozone. Mice were exposed to ozone for four days and harvested on day 7 as depicted in Figure 4.

**Supplementary Figure 6.**
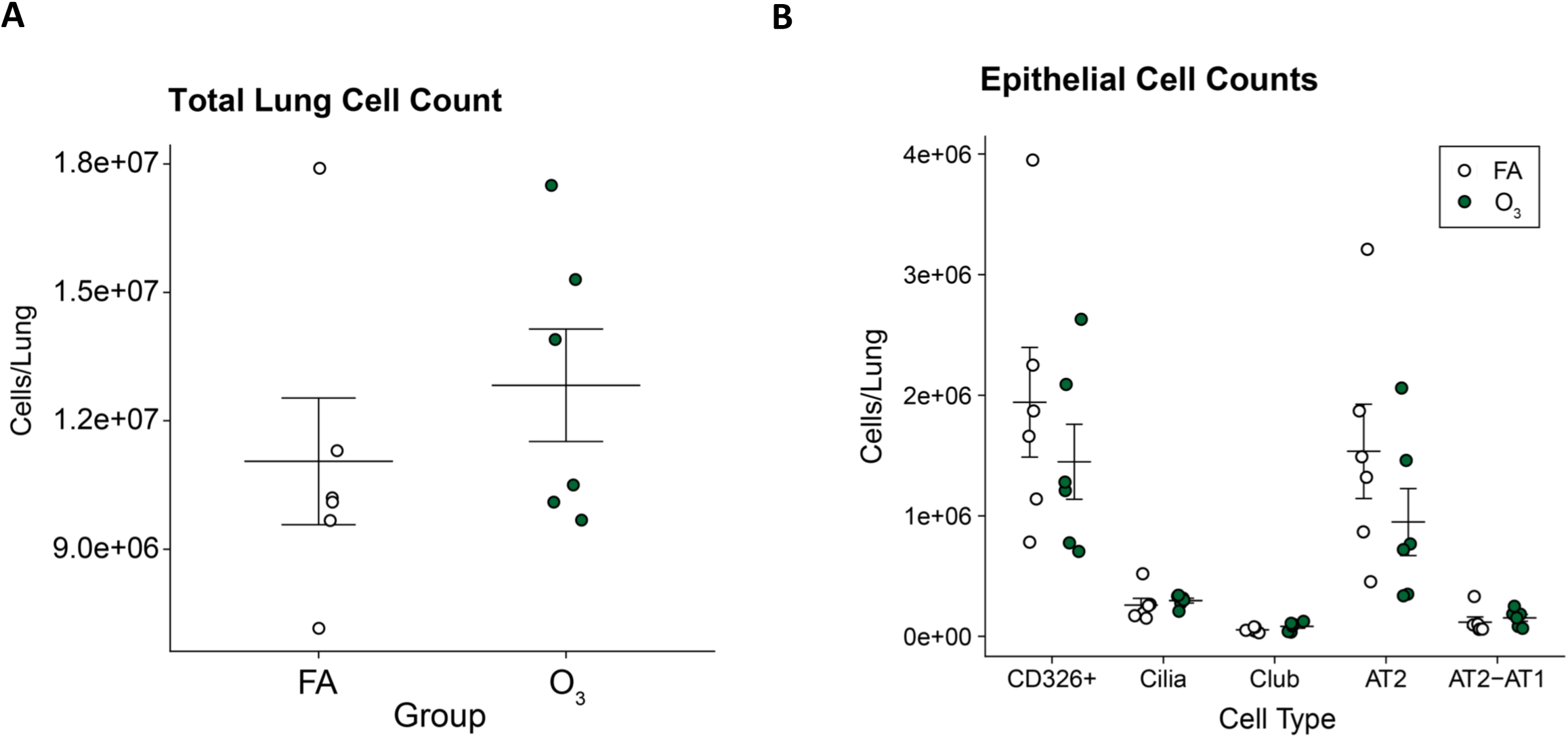
Cell counts of mice exposed to ozone for four days and harvested on day 7. **A.** Total cell counts after lung digested determined manually with a hemacytometer. **B.** Epithelial cell counts determined by flow cytometry.

**Supplementary Figure 7.**
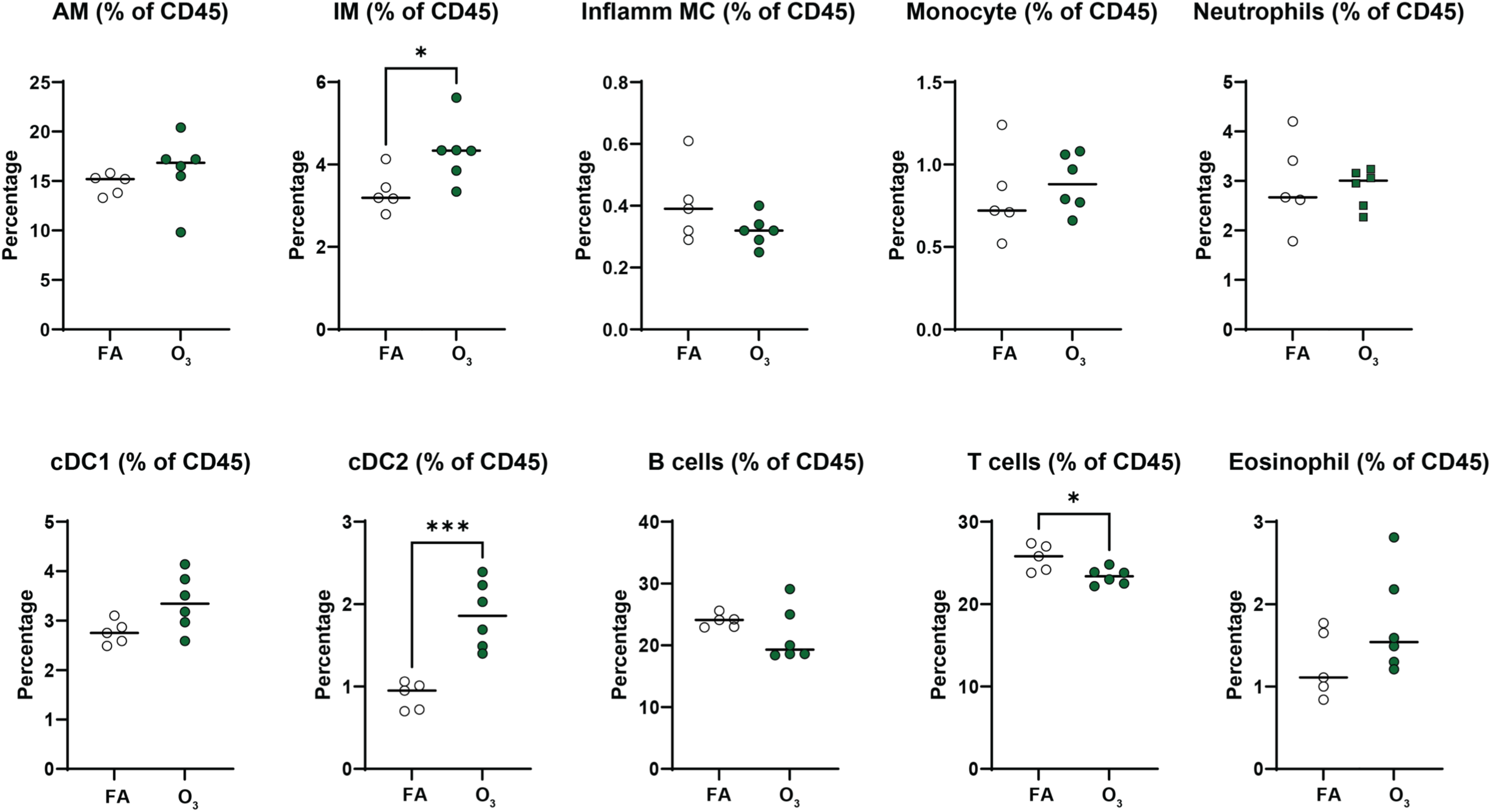
Frequencies of CD45+ leukocytes determined by flow cytometry.

**Supplementary Figure 8.**
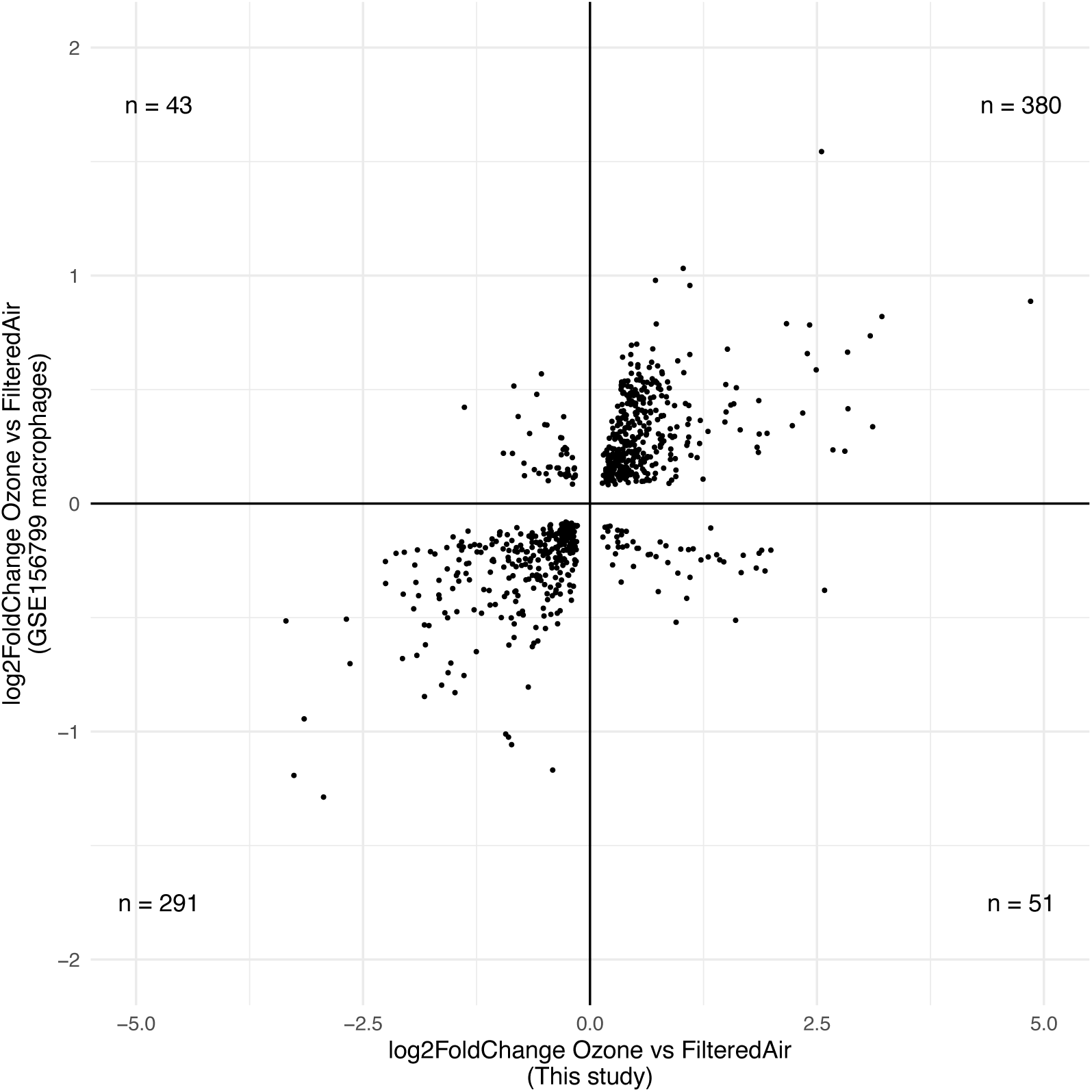
Comparison of AM DEGs in this study vs. GSE156799. Shrunken log2 fold change values for DEGs detected in each study are shown.

**Supplementary Figure 9.**
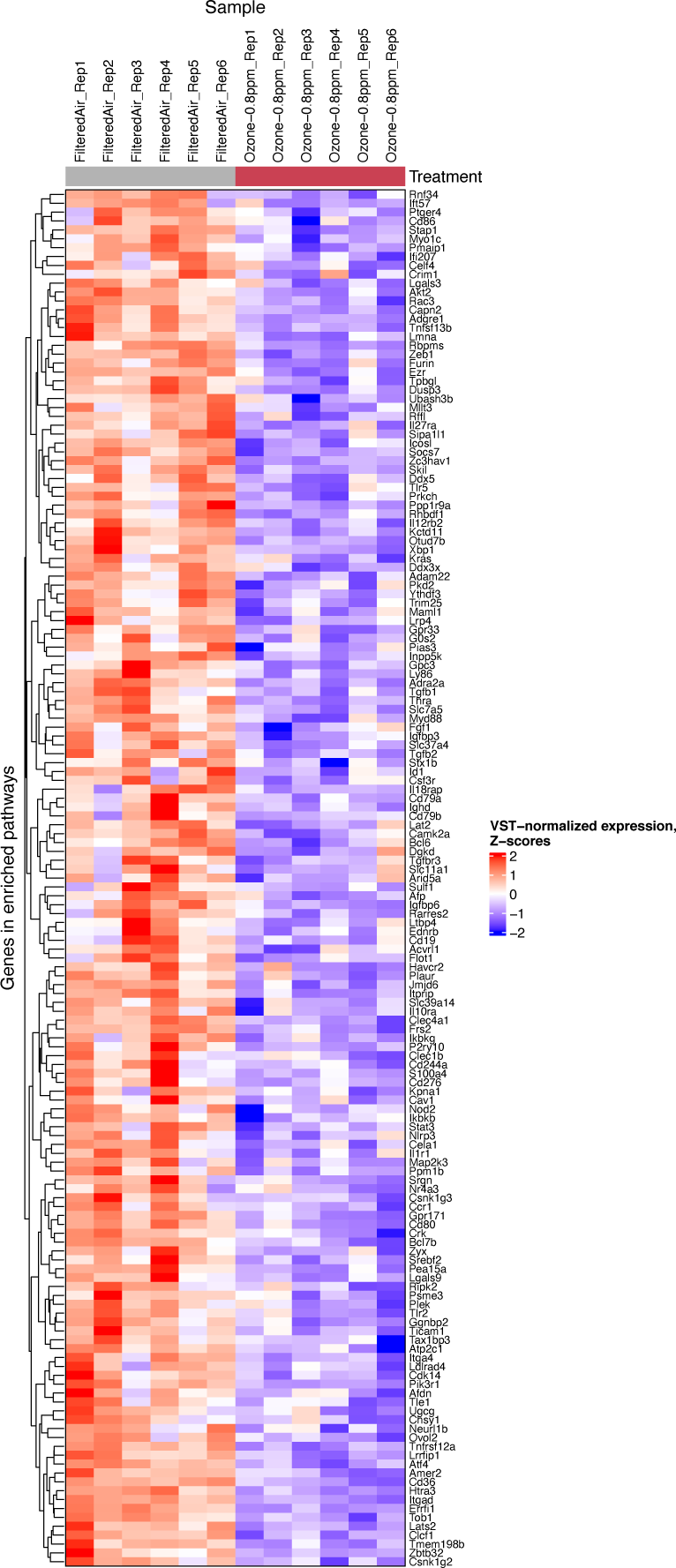
Expression of genes corresponding to pathways shown in Figure 8B.

