## Supplementary Table 1 for "Alveolar macrophages play a key role in tolerance to ozone"

**Supplementary Table 1. Antibodies used in flow cytometry experiments**

|  | **Antibody** | **Clone** | **Company** | **Catalog #** |
| --- | --- | --- | --- | --- |
| Epithelial cell flow cytometry | CD16/C32 (FcγRIII/FcγRII) | 2.4G2 | BD Biosciences | 553141 |
|  | CD24 PerCP-Cy5.5 | M1/69 | BioLegend | 101824 |
|  | MHCII Alexa 700 | M5/114.15.2 | BioLegend | 107618 |
|  | Ki67 PE | B56 | BD Biosciences | 556027 |
|  | CD326 PE-Dazzle | G8.8 | BioLegend | 118236 |
|  | Zombie NIR | - | BioLegend | 423106 |
|  | CD31 PB (BV421) | 390 | BioLegend | 102422 |
|  | T1 alpha biotin | 8.1.1 | BioLegend | 127404 |
|  | Streptavidin BV605 | - | BioLegend | 405229 |
|  | CD45 BV785 | 30-F11 | Biolegend | 103149 |
|  | CD104 APC | 346-11A | Biolegend | 123612 |
| Lung leukocyte flow cytometry | CD16/C32 (FcγRIII/FcγRII) | 2.4G2 | BD Biosciences | 553141 |
|  | CD11b BV605 | M1/70 | Biolegend | 101257 |
|  | CD11c PerCP/Cy5.5 | N418 | Biolegend | 117328 |
|  | CD88 PE/Cy7 | 20/70 | Biolegend | 135810 |
|  | CD103 PE/Dazzle 594 | 2E7 | Biolegend | 121429 |
|  | Siglec-F PE | E50-2440 | BD Biosciences | 552126 |
|  | CD3e Biotin | 145-2C11 | Biolegend | 100304 |
|  | CD19 Biotin | 6D5 | Biolegend | 115504 |
|  | Ly-6C APC/Cy7 | AL-21 | Biolegend | 560596 |
|  | Ly-6G FITC | 1A8 | Biolegend | 127605 |
|  | I-A/E Pacific Blue | M5/114.15.2 | Biolegend | 107620 |
|  | Zombie Aqua | - | BioLegend | 423101 |
