## Supplementary Table 2 for "Alveolar macrophages play a key role in tolerance to ozone"

**Supplementary Table 2. Primer sequences used for qRT-PCR.**

| **Gene** | **Forward Primer** | **Reverse Primer** |
| --- | --- | --- |
| *Alox15* | CTCTCAAGGCCTGTTCAGGA | GTCCATTGTCCCCAGAACCT |
| *Alox5* | CTGCTGGACAAGGCATTCTA | TCCATCCCTCAGGACAATCT |
| *Cat* | CAAGTTGGTTAATGCAGATGGAG | ATCTTCCTGAGCAAGCCTTC |
| *Gclc* | ACATCTACCACGCAGTCAAG | ACATCGCCTCCATTCAGTAAC |
| *Gclm* | ACAAGACACAGTTGGAGCAG | GTGAGTCAGTAGCTGTATGTCA |
| *Gpx1* | GACTACACCGAGATGAACGAT | AGGTCGGACGTACTTGAGG |
| *Hmox1* | ACACTCTGGAGATGACACCT | TTGTGTTCCTCTGTCAGCATC |
| *Nfe2l2* | TGATGGACTTGGAGTTGCC | TCAAACACTTCTCGACTTACTCC |
| *Nqo1* | GCCAATGCTGTAAACCAGTTG | GCTCCATGTACTCTCTTCAGG |
| *Rps20** | CGATTTATTCAGTTGTCTTAGGCATC | AGTCCTTCTGAGATTGTTAAGCAG |
| *Sod1* | GTCCTTTCCAGCAGTCACAT | GGTTCCACGTCCATCAGTATG |
| *Sod2* | TGTCAGCTTCTCCTTAAACTTCT | GGACAAACCTGAGCCCTAAG |

*gene used for normalization
